# Representing Multiple Visual Objects in the Human Brain and Convolutional Neural Networks

**DOI:** 10.1101/2023.02.28.530472

**Authors:** Viola Mocz, Su Keun Jeong, Marvin Chun, Yaoda Xu

**Affiliations:** Visual Cognitive Neuroscience Lab, Department of Psychology, Yale University, New Haven, CT 06520, USA; Department of Neuroscience, Yale School of Medicine, New Haven, CT 06520, USA; Department of Psychology, Chungbuk National University, South Korea

## Abstract

Objects in the real world often appear with other objects. To recover the identity of an object whether or not other objects are encoded concurrently, in primate object-processing regions, neural responses to an object pair have been shown to be well approximated by the average responses to each constituent object shown alone, indicating the whole is equal to the average of its parts. This is present at the single unit level in the slope of response amplitudes of macaque IT neurons to paired and single objects, and at the population level in response patterns of fMRI voxels in human ventral object processing regions (e.g., LO). Here we show that averaging exists in both single fMRI voxels and voxel population responses in human LO, with better averaging in single voxels leading to better averaging in fMRI response patterns, demonstrating a close correspondence of averaging at the fMRI unit and population levels. To understand if a similar averaging mechanism exists in convolutional neural networks (CNNs) pretrained for object classification, we examined five CNNs with varying architecture, depth and the presence/absence of recurrent processing. We observed averaging at the CNN unit level but rarely at the population level, with CNN unit response distribution in most cases did not resemble human LO or macaque IT responses. The whole is thus not equal to the average of its parts in CNNs, potentially rendering the individual objects in a pair less accessible in CNNs during visual processing than they are in the human brain.

## Introduction

In everyday visual perception, objects are rarely encoded in isolation, but often with other objects appearing in the same scene. It is thus critical for primate vision to recover the identity of an object regardless of whether or not other objects are encoded concurrently. Both monkey neurophysiological and human fMRI studies have reported the existence of averaging for representing multiple objects at high-level vision. Specifically, in macaque inferotemporal (IT) cortex that is not category selective, neuronal response amplitude to a pair of unrelated objects can be approximated by the average response amplitude of each object shown alone (Zoccolan et al., 2005); similarly, in human occipital-temporal cortex, fMRI response pattern to a pair of unrelated objects can be predicted by the average fMRI response pattern of each object shown alone (MacEvoy & Epstein, 2009, 2011; Reddy & Kanwisher, 2007; Reddy, et al., 2009; Jeong & Xu, 2017). The whole is thus equal to the average of parts at high-level primate vision (note that responses in category-selective regions may exhibit both the averaging and winner-take-all responses depending on the stimuli shown, see Reddy & Kanwisher, 2007; Reddy et al., 2009; Bao & Tsao, 2018; Kliger & Yovel, 2020). Such a representational scheme can effectively avoid response saturation, especially for neurons responding vigorously to each constituent object, and prevent the loss of identity information when objects are encoded together (MacEvoy & Epstein, 2009). Such tolerance to the encoding context, together with the ability of the primate high-level vision to extract object identity across changes in other non-identity features (such as viewpoint, position and size), has been argued as one of the hallmarks of primate high-level vision that allows us to rapidly recognize an object under different viewing conditions (DiCarlo and Cox, 2007; DiCarlo et al., 2012; Tacchetti et al., 2018).

In neurophysiological studies, averaging was assessed at the single unit level by documenting the slope of single neuron response amplitudes to an object pair and the constituent objects. In fMRI studies, however, averaging was measured at the population level by correlating the voxel response patterns. This raises the intriguing question of whether averaging can be directly seen at the single unit level in the slope of single fMRI voxel response amplitudes. Given that each fMRI voxel contains around a million neurons and that fMRI is an indirect measure of neuronal activity (Huettel et al., 2009), it is not obvious that single neuron response properties are directly observable in single fMRI voxels. Nevertheless, monkey IT neurons are organized into clusters of 0.5 mm in diameter containing neurons with similar tuning profiles (e.g., Wang et al., 1996; Tsunoda et al., 2001). This is a meso-scale organization matching the resolution of fMRI. Such functional smoothness in neuronal representation has been argued to enable the success of fMRI to capture brain functions (Guest & Love, 2017). The first goal of this study is therefore to test if averaging in individual voxels indeed exists. If it does, we will further test if voxels showing better averaging in response amplitude may exhibit better response pattern averaging at the population level. Although it is reasonable to assume that averaging in fMRI response patterns and in the individual voxels correspond to the same underlying neural mechanism, this assumption, however, has not been directly tested. Here we will examine fMRI response patterns and individual voxel responses from human lateral occipital cortex (LO), a higher ventral visual object processing region homologous to the macaque IT, to gain a better understanding of the relationship between unit and population responses in predicting paired object responses from those of single objects.

Convolutional neural networks (CNNs) are currently considered as one of the best models of primate vision, achieving human-level performance in object recognition tasks and showing correspondences in object representation with the primate ventral visual system (e.g., Khaligh-Razavi & Kriegeskorte, 2014; Cichy et al., 2016; Xu & Vaziri-Pashkam, 2021a). Meanwhile, there still exist large discrepancies in visual representation and performance between the CNNs and the primate brain (Serre, 2019), with CNNs only able to account for about 60% of the representational variance seen in primate high-level vision (Yamins, et al., 2014; Bao et al., 2020; Kar et al., 2019; Xu & Vaziri-Pashkam, 2021a & 2021b). While CNNs are fully image computable and accessible, they are also “blackboxes” - extremely complex models with millions or even hundreds of millions of free parameters whose general operating principles at the algorithmic level (Marr, 1982) remain poorly understood (e.g., Kay, 2018).

Because CNNs are trained with natural images containing a single target object appearing in a natural scene, it is unclear that objects are represented as distinctive units of visual processing and that an averaging relationship for representing multiple objects would automatically emerge. Moreover, neural averaging is similar to divisive normalization previously proposed to explain attentional effects in early visual areas (Carandini & Heeger, 2012; Reynolds & Heeger, 2009; Heeger, 1992). Such a normalization process involves dividing the response of a neuron by a factor that includes a weighted sum of the activity of a pool of neurons through feedforward, lateral or feedback connections. Given that some of the well-known CNNs have no lateral or feedback connections, such as Alexnet (Krizhevsky et al., 2012), VGG-16 (Simonyan & Zisserman, 2014), Googlenet (Szegedy et al., 2015) and Resnet-50 (He et al., 2016), these CNNs should not be expected to show response averaging for representing multiple objects. Nonetheless, by assessing the slope across CNN unit responses, Jacob et al. (2021) reported that higher layers of VGG16 exhibited averaging similar to that of macaque IT neurons. This is a rather unexpected finding. Because the slopes across all CNN units were averaged, it is unclear whether individual units indeed exhibit responses similar to those found in the primate brain. Likewise, it is presently unknown whether averaging at the CNN unit level would lead to averaging at the response pattern level. Answers to the above questions are critical if we want to better understand the nature of visual object representations in CNNs and whether CNNs represent visual objects similarly as the primate brain. To do so, here we examined single unit and population responses from the higher layers of four CNNs pretrained for object categorization with varying architecture and depth. We additionally examined the higher layers of a CNN with recurrent processing to test whether averaging at both the single unit and population level may emerge when feedback connections are present in a network. As with the fMRI data, we also examined the relationship between unit and population responses by testing whether units showing better averaging in response amplitude would exhibit better pattern averaging at the population level.

## Results

In this study, we analyzed a previous fMRI data set (Jeong & Xu, 2017) where participants viewed blocks of images showing either single object or object pairs selected from four different object categories (bicycle, couch, guitar, or shoe) and performed a 1-back repetition detection task (Figure 1A). We selected the 75 most reliable voxels from LO in our analysis to equate the number of voxels from each participant and to increase power (Tarhan & Konkle, 2019). However, all results remained virtually identical when we included all voxels from LO. This indicates the stability and robustness of the results which did not depend on including the most reliable voxels.

**Figure 1.**
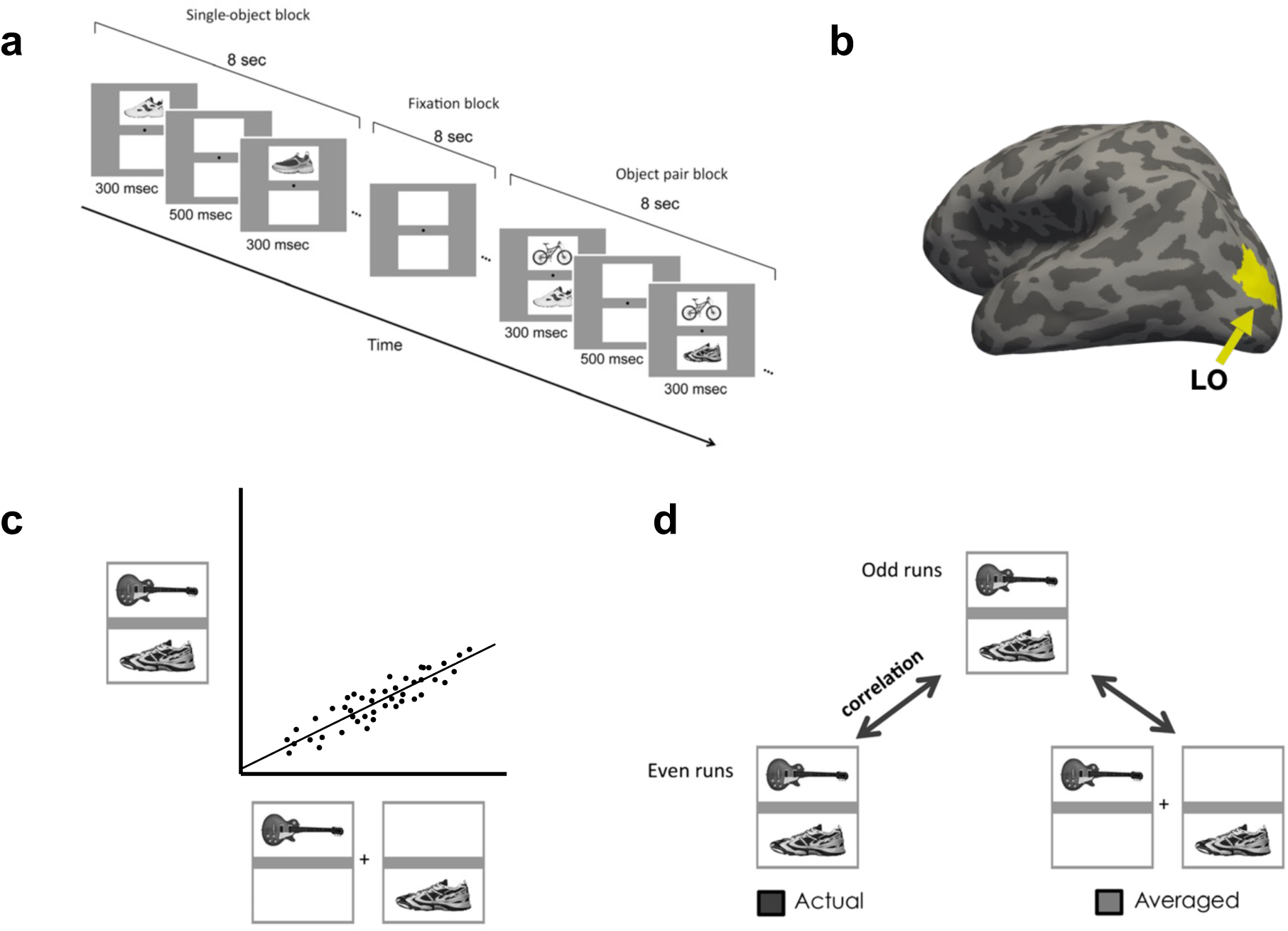
Experimental and analysis details. **a.** Example trials of the main experiment. In single-object blocks, participants viewed a sequential presentation of single objects (different exemplars) from the same category at a fixed spatial location. In object pair blocks, they viewed a sequential presentation of object pairs from two different categories. The location of the objects from each of the two categories was fixed within a block. The task was to detect a 1-back repetition of the same object. In object pair blocks, the repetition could occur in either object category. The repetition occurred twice in each block. **b.** Inflated brain surface from a representative participant showing LO. **c.** A schematic illustration of the unit response analysis. We extracted, across all the included fMRI voxels/CNN units, the slope of the linear regression between an fMRI voxel/CNN unit’s response to a pair of objects and its summed response to the corresponding single objects shown in isolation. **d.** A schematic illustration of the population response analysis. For LO, an object pair was correlated with itself across the odd and even halves of the runs (actual) and with the average of its constituent objects shown in isolation across the odd and even halves of the runs (average). For CNNs, because there is no noise, the object pair was simply correlated with the average of its constituent objects in isolation.

We also analyzed the higher layers from five CNNs pre-trained using ImageNet images (Deng et al., 2009) to perform object categorization. The CNNs examined included both shallower networks, Alexnet (Krizhevsky et al., 2012) and VGG-19 (Simonyan & Zisserman, 2014), and deeper networks, Googlenet (Szegedy et al., 2015) and Resnet-50 (He et al., 2016). We also included a recurrent network, Cornet-S, that has been shown to capture the recurrent processing in macaque IT cortex with a shallower structure and argued to be one of the current best models of the primate ventral visual system (Kar et al., 2019; Kubilius et al., 2019). We further analyzed three versions of Resnet-50 trained on stylized versions of ImageNet images (Geirhos et al., 2019) to examine how reducing texture bias and increasing object shape processing in a CNN would impact its representation of multiple objects. For all CNNs, we created a comparable set of activation patterns as the brain data. In the fMRI experiment, objects were presented within a white rectangle on a gray background. It is possible that this gray background could affect object averaging as CNNs wouldn’t “filter out” the gray background as human participants would. We thus examined CNN responses to objects on both the gray and white backgrounds.

### Evaluating the Unit Response to Single and Paired Objects

In this analysis, we extracted, across all the included fMRI voxels/CNN units, the slope of the linear regression between an fMRI voxel/CNN unit’s response to a pair of objects and its summed response to the corresponding single objects shown in isolation. We averaged the slopes across all the human participants. The average slope should be 0.5 if the single fMRI voxel/CNN unit response to an object pair can be perfectly predicted by the average response of the corresponding objects shown alone. In a previous single-cell analysis of monkey IT, a slope of about 0.55 was reported (Zoccolan et al., 2005). We thus compared our slope results to both 0.5 and 0.55 as baselines. Following Jacob et al. (2021), we only selected CNN units that showed non-zero responses to objects at both locations.

#### Human LO

The average slope for all participants when including the 75 most reliable voxels and all the voxels were 0.573 and 0.561, respectively (Figure 3A). These average slopes did not differ from the reported IT neuron slope of 0.55 (*t(9)* = 0.866, *p* = .694, *d* = 0.273, for the 75 most reliable voxels; and *t(9*) = 0.406, *p* = .694, *d* = 0.128, for all the voxels; corrected for multiple comparisons using the Benjamini-Hochberg method for two comparisons, see Benjamini & Hochberg, 1995). These slopes, however, did deviate from the perfect averaging slope of 0.5 (*t(9)* = 2.741, *p* = .046, *d* = 0.867, for the 75 most reliable voxels; and *t(9)* = 2.22, *p* = .054, *d* = 0.701 for all the voxels; corrected). Inspection of the distribution of the voxel response aggregated across all participants revealed a distribution around the line of best fit, similar to a normal distribution and the distribution seen in the IT neuron responses (Figure 2A). Overall, we observed averaging in single fMRI voxels in human LO comparable to that of single neurons in macaque IT.

**Figure 2.**
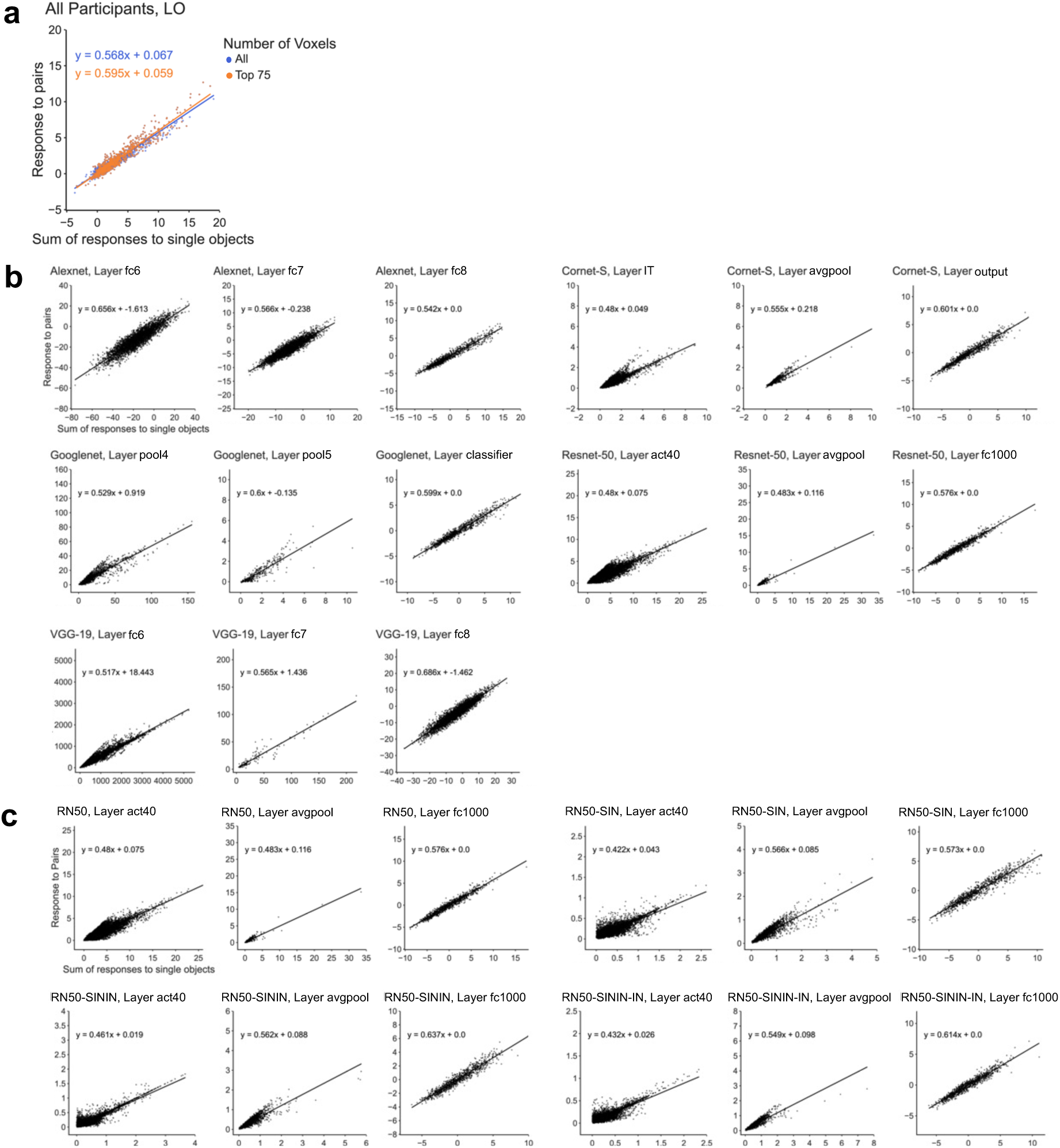
Single unit response amplitude distribution results. **a.** Single voxel response to object pairs plotted against the sum of the individual object responses in LO for all of the participants for both the top 75 most reliable voxels and all the voxels. Units are in beta weights. **b.** Responses of the five CNNs (pretrained on the original ImageNet images) to paired and single objects on gray background. **c.** Responses of Resnet-50 to paired and single objects on gray background. Resnet-50 was pretrained either with the original ImageNet images (RN50-IN), the stylized ImageNet Images (RN50-SIN), both the original and the stylized ImageNet Images (RN50-SININ), or both sets of images, and then fine-tuned with the stylized ImageNet images (RN50-SININ-IN).

**Figure 3.**
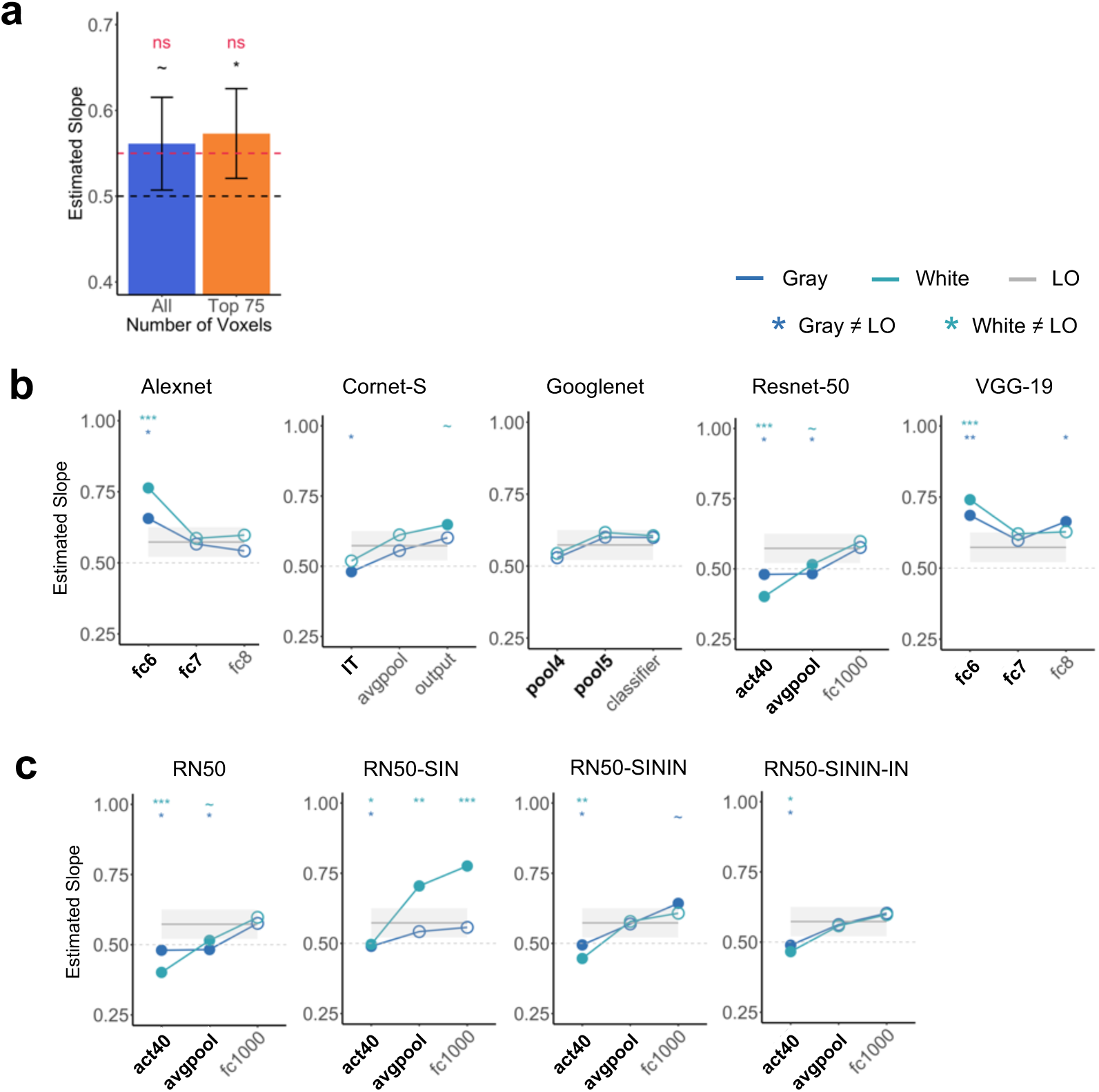
Single unit response summary results. **a.** The averaged slope across all participants for both the top 75 most reliable LO voxels and for all the LO voxels. The average slope is compared to the slope of a perfect averaging (0.5) as well as the slope reported in single cells in macaque IT (0.55). **b.** Comparing LO slope with those of 5 CNNs with the objects appearing on both the gray and white backgrounds. Bold layers were those that showed best correspondence with LO as shown by Xu & Vaziri-Pashkam (2021a). For each layer, significant values for pairwise t tests against the slope of LO are marked with asterisks at the top of the plot. **c.** Comparing LO slope with those of Resnet-50 under different training regimes. Resnet-50 was pretrained either with the original ImageNet images (RN50-IN), the stylized ImageNet Images (RN50-SIN), both the original and the stylized ImageNet Images (RN50-SININ), or both sets of images, and then fine-tuned with the stylized ImageNet images (RN50-SININ-IN). All t tests were corrected for multiple comparisons using the Benjamini-Hochberg method for 2 comparisons in LO, and for 6 comparisons (3 layers x 2 background colors) for each CNN. Error bars and ribbons represent the between-subjects 95% confidence interval of the mean. ∼ .05 < p < .10, * p <.05, ** p <.01, *** p <.001.

#### CNNs

When the slopes from the sampled CNN layers were directly tested against the slope obtained from the human LO for the 75 most reliable voxels, 8 out of the 15 (53%) examined layers did not differ from LO for either the white or gray background images (*ts* < 2.05, *ps* > .11; all others, *ts* > 2.82, *ps* < .06; see the asterisks marking the significance levels on Figure 3B; all pairwise comparisons reported here and below were corrected for multiple comparisons using the Benjamini-Hochberg method). When we further examined the layers that showed the greatest correspondence to LO as reported in a previous study (Xu & Vaziri-Pashkam, 2021a), 4 out of the 9 (44%) layers were not significantly different from LO (see the layers marked with bold font in Figure 3B). There was little effect of image background (i.e., whether the images appeared on the white or gray backgrounds), with only 3 out the 15 (20%) layers showing a discrepancy when comparing with LO (i.e., with performance on one background being similar and performance on the other background being different from that of LO). For the other 12 layers, performance (as compared to LO) did not differ across the two backgrounds. Overall, among the CNNs tested, Googlenet best resembled the human LO, with all of its sampled layers showing no significant difference from LO. Interestingly, the recurrent network examined, Cornet-S, did not seem to behave differently from the other networks.

In Jacob et al. (2021), there was a noticeable improvement in unit averaging from lower to higher layers. Here we found that, except for Googlenet, the other CNNs all showed an improvement between the first two examined layers. Resnet-50 additionally showed an improvement between the last two examined higher layers.

Close inspection revealed that the distribution of the CNN unit responses to object pair and single objects varied greatly across the examined CNN layers, with some showing a distribution resembling that of a normal distribution and those of LO voxels and IT neurons, but others deviating greatly from such a distribution even when the average slope was close to 0.5 or 0.55 (Figure 2B). For example, although layer fc1000 in Resnet-50 and layer pool4 in Googlenet had slopes close to 0.5 and 0.55 (0.576 and 0.529, respectively), Resnet-50 had an approximately normal distribution while Googlenet does not. Additionally, although layer fc1000 in Resnet-50 and layer fc6 in Alexnet had approximately normal distributions, their slopes were very different from each other (0.576 and 0.656, respectively). This suggested that the representation of multiple objects in some higher CNN layers was different from that of the human LO and macaque IT.

### Evaluating the Pattern Response to Single and Paired Objects

In this analysis, we wanted to replicate the existence of averaging in fMRI response patterns as reported in Jeong and Xu (2017) such that response patterns for an object pair may be predicted by the average pattern of the constituent objects shown in isolation. We also tested whether such averaging existed in CNN unit response patterns. To analyze the fMRI pattern responses, as in Jeong and Xu (2017), we conducted a split-half analysis to account for measurement noise (as correlation could never reach 1 even for the same condition across different halves of the data due to the presence of measurement noise in fMRI data). Specifically, we divided the runs into even and odd halves and averaged the response patterns for each condition within each half of the runs. We then correlated the response patterns of the same object pairs across even and odd halves (actual pair correlation) and correlated the response patterns of an object pair from one half and the average response pattern of the constituent single objects in the other half (average pair correlation). Finally, using the actual pair correlation as a baseline, we derived a noise-normalized correlation by dividing the average pair correlation by the actual pair correlation. If averaging existed in response patterns, then the response pattern of an object pair would be as similar to itself in the other half of the runs as it would be to the averaged response patterns of the single objects in the other half of the runs. We would then have a noise-normalized correlation no different from 1. For the CNNs, because there was no noise, we simply correlated the response pattern of a pair of objects with the average response pattern of the constituent single objects. We then compared this correlation to that of LO. We additionally assessed whether a weighted average of single objects would improve pattern averaging. We did so by systematically varying the contribution of the two single objects’ response patterns from 0% to 100%, with 10% increments in between.

#### Human LO

In LO, the noise-normalized correlation did not differ from 1 for neither the Top 75 voxels (*t(9)* = 1.588, *p* = .146, *d* = 0.502; one-tailed as only comparison in one direction was meaningful here) nor all voxels (*t(9)* = 0.498, *p* = .315, *d = 0.157*; one-tailed) (corrected for two comparisons) (Figure 4A). Additional analyses showed that weighing the two constituent objects equally yielded the best averaging prediction (Figure 5C). These results replicated those of Jeong & Xu (2017) and showed the existence of averaging in fMRI response patterns in human LO.

**Figure 4.**
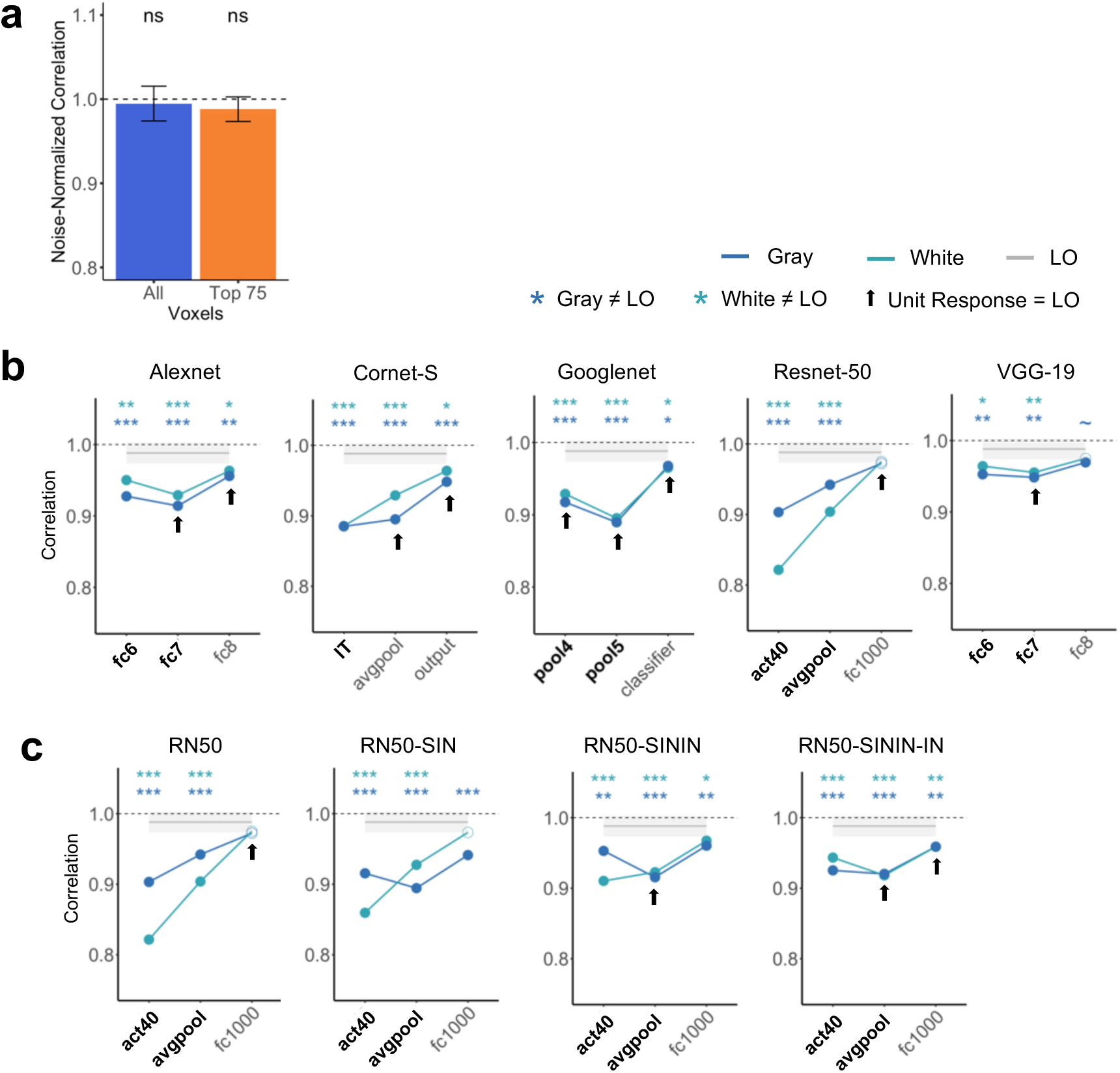
Pattern response results. **a.** Noise-normalized correlation between the pattern response to an object pair and the average pattern response to the individual objects in LO for the top 75 most reliable voxels and all the voxels. A one-sided t-test was conducted comparing the correlations to 1. **b.** Comparing pattern averaging in LO with those of five CNNs with the objects appearing on both the gray and white backgrounds. Bold layers were those that showed best correspondence with LO as shown by Xu & Vaziri-Pashkam (2021a). For each layer, significant values for pairwise t tests against LO are marked with asterisks at the top of the plot. Layers showing comparable averaging in unit response amplitude to that of LO are marked with an arrow. **c.** Comparing pattern averaging in LO with that of Resnet-50 under different training regimes. Resnet-50 was pretrained either with the original ImageNet images (RN50-IN), the stylized ImageNet Images (RN50-SIN), both the original and the stylized ImageNet Images (RN50-SININ), or both sets of images, and then fine-tuned with the stylized ImageNet images (RN50-SININ-IN). All t tests were corrected for multiple comparisons using the Benjamini-Hochberg method for 2 comparisons in LO, and for 6 comparisons (3 layers x 2 background colors) for each CNN. Error bars and ribbons represent the between-subjects 95% confidence interval of the mean. ∼ .05 < p < .10, * p <.05, ** p <.01, *** p <.001.

**Figure 5.**
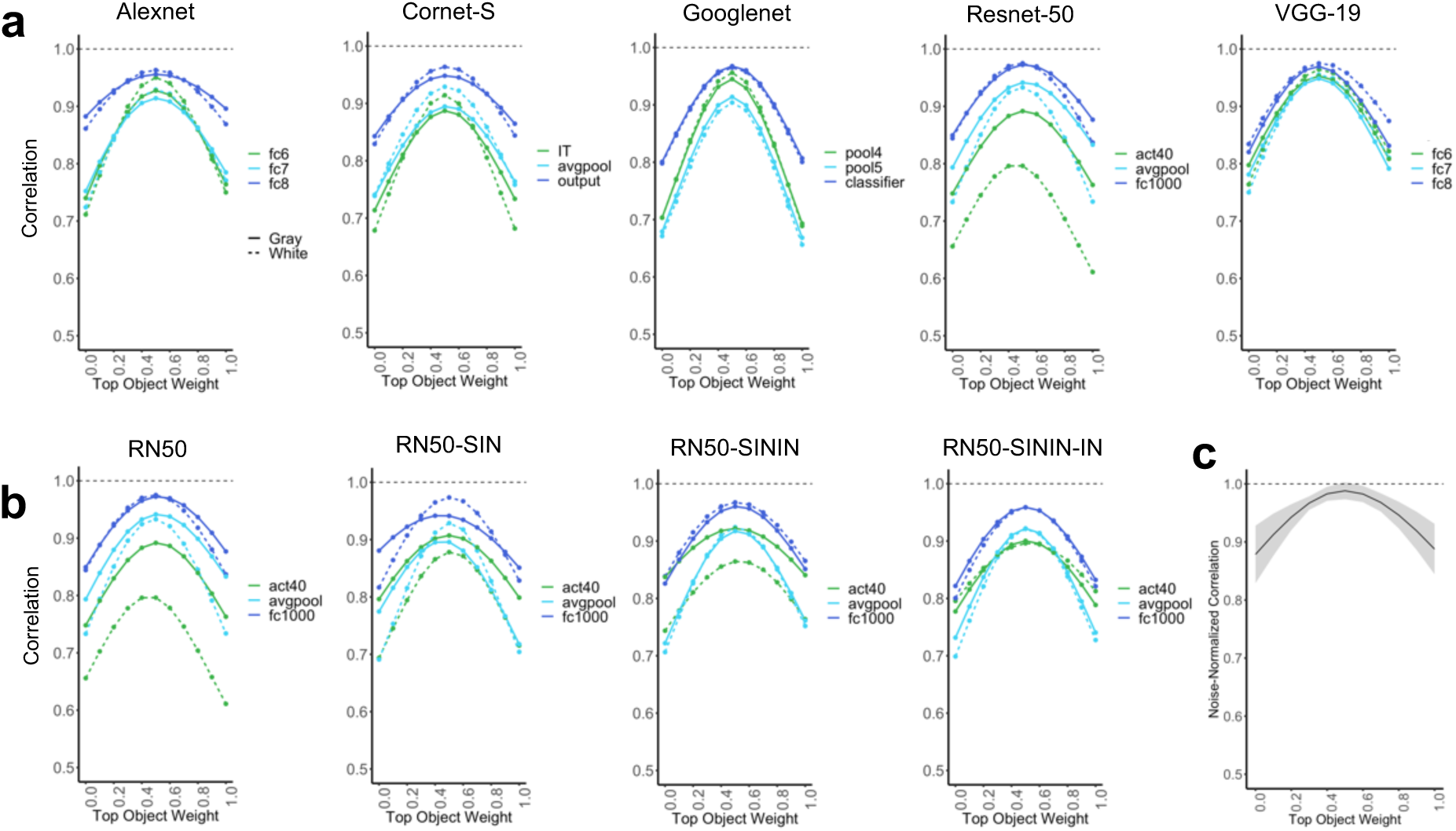
Pattern averaging results with a weighted average of the two single objects, where the weight of each object varies between 0 and 1 with increments of 0.1. **a.** Weighted pattern averaging for the five CNNs trained on the original ImageNet images. **b.** Weighted pattern averaging for Resnet-50. Resnet-50 was pretrained either with the original ImageNet images (RN50-IN), the stylized ImageNet Images (RN50-SIN), both the original and the stylized ImageNet Images (RN50-SININ), or both sets of images, and then fine-tuned with the stylized ImageNet images (RN50-SININ-IN). **c.** Weighted pattern averaging for LO including all the voxels.

#### CNNs

Overall, all sampled CNN layers showed a correlation in pattern response greater than .8. However, with the exception of the last sampled layer in Resnet-50, for all others, the correlation was significantly less than the noise-normalized correlation of the top 75 voxels in LO regardless of the background color (see the asterisks marking the significance levels in Figure 4B). This suggests that pattern averaging of single object responses could not fully predict that of the paired objects in most of the sampled CNN layers, different from the results obtained in human LO (Figure 4B). This was true even for layers showing strong averaging in single unit response amplitudes as shown in Figures 3B and marked on Figure 4B, indicating a discrepancy between unit and pattern responses in these CNN layers. This was also true for Cornet-S, the recurrent network that included feedback connections among the units to better model the primate visual system. Of all the CNN layers sampled, only the final layer of Resnet-50 exhibited pattern correlation and single unit responses matching those of LO (see Figures 3B and 34). That being said, both Cornet-S and Resnet-50 showed an improvement in averaging between the first two sampled layers, and all CNNs showed an improvement between the last two sampled layers, although such improvement was still not sufficient to reach the performance of LO. Additional analyses showed that, similar to LO, an equally weighted averaging of both constituent objects yielded the best pattern prediction (Figure 5A).

### Evaluating the Relationship between Unit Response and Pattern Response

In this analysis, we wanted to gain a better understanding of the relationship between unit and population responses and why there might be a discrepancy between unit and pattern responses in CNNs. We evaluated whether single units showing better averaging in response amplitude also show better averaging in response pattern. We divided LO voxels/CNN units into two groups by whether the slope in response amplitude averaging for the single voxel/CNN unit was near or far from 0.5, with the *near* condition defined as having a slope between 0.45 and 0.55 and the *far* condition defined as having a slope less than 0.45 or greater than 0.55. We then assessed averaging in response patterns in the near and far groups of voxels and units separately. We also noted the percentage of voxels/CNN units in the two groups.

#### Human LO

We included all LO voxels in this analysis. Overall, approximately 75% of the total voxels across all of the participants had a slope near 0.5 (Figure 6A). Such a distribution is consistent with the voxel distribution seen in Figure 6A. While there was no difference between the noise-normalized pattern correlation and 1 for the near voxels (*t(9)* = 1.942, *p* = .958, *d* = 0.614; corrected), the difference was marginally significant for the far voxels (*t(9)* = 2.029, *p* = .073, *d* = 0.642; corrected), with a significant difference between the near and far voxels (*t(9)* = 2.52, *p* = .033, *d* =1.06). In human LO, voxels with a better averaging response in response amplitude thus also showed better averaging in response patterns.

**Figure 6.**
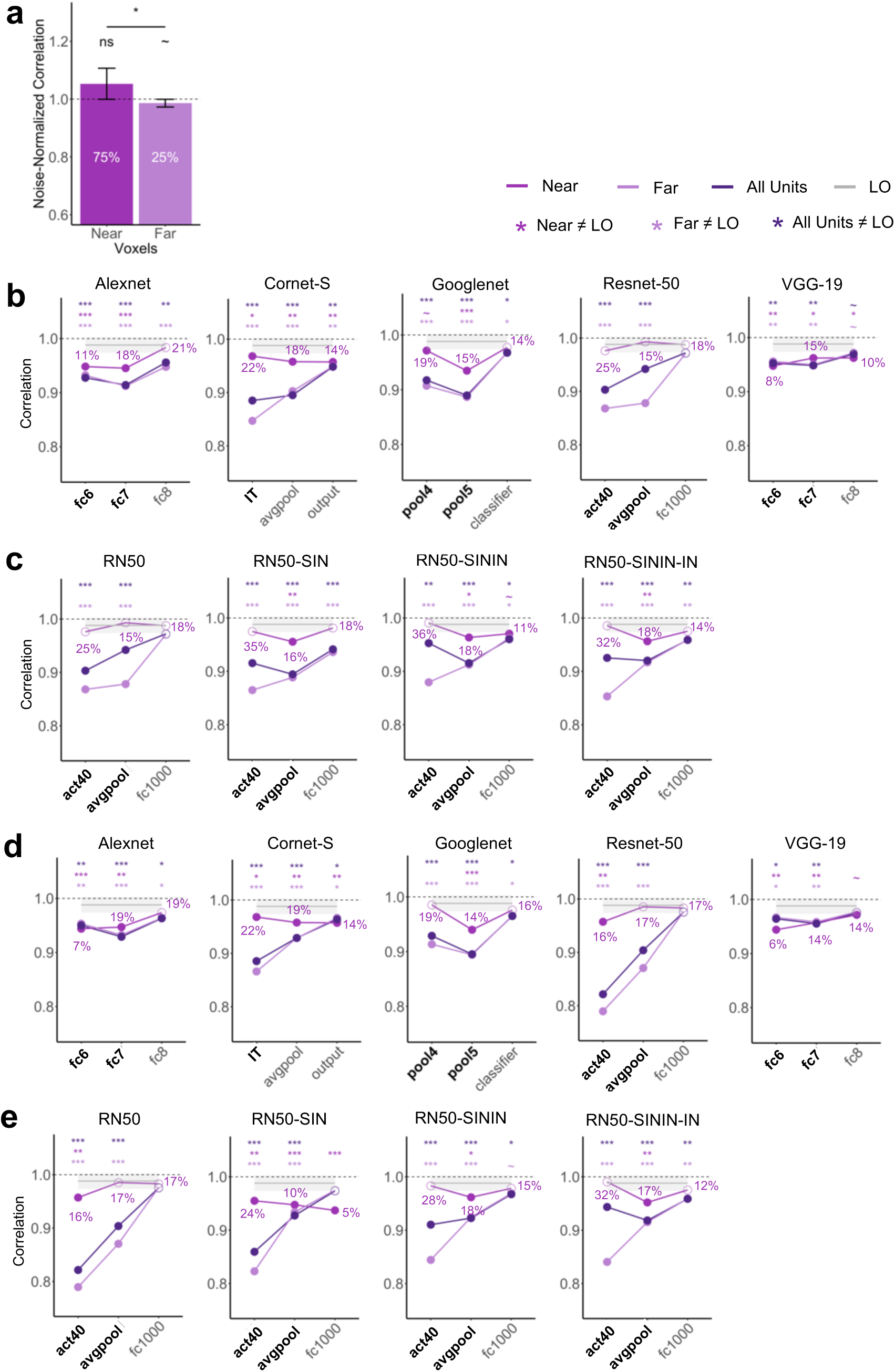
Relationship between unit and pattern responses. **a.** Noise-normalized correlation between the pattern response to the pair and the average pattern response to the individual objects in LO for voxels that have slope near 0.5 (0.45 < slope < 0.55) and for voxels that have slope far from 0.5 (slope < 0.45 or slope > 0.55). The percentage of voxels considered to be near or far from slope 0.5 are marked. A one-sided t-test was conducted comparing the correlations to 1. **b.** CNN pattern averaging results for images with gray background, separately for CNN units with slope near 0.5, units with slope far from 0.5, and all units, all compared with pattern averaging from LO. The percentages of units near slope 0.5 are marked. Bold layers were those that showed best correspondence with LO as shown by Xu & Vaziri-Pashkam (2021a). For each layer, significant values for pairwise t tests against the slope of LO are marked with asterisks at the top of the plot. **c.** For Resnet-50 under different training regimes, CNN pattern averaging results for images with gray background, separately for CNN units with slope near 0.5, units with slope far from 0.5, and all units, all compared with pattern averaging from LO. Resnet-50 was pretrained either with the original ImageNet images (RN50-IN), the stylized ImageNet Images (RN50-SIN), both the original and the stylized ImageNet Images (RN50-SININ), or both sets of images, and then fine-tuned with the stylized ImageNet images (RN50-SININ-IN). **d.** CNN pattern averaging results for images with white background, separately for the three groups of units as in b. **e.** Resnet-50 pattern averaging results for images with white background, separately for the three groups of units as in c. All t tests were corrected for multiple comparisons using the Benjamini-Hochberg method for 2 comparisons in LO, and for 9 comparisons (3 layers x 3 groups of units) in each CNN. Error bars and ribbons represent the between-subjects 95% confidence interval of the mean. ∼ .05 < p < .10, * p <.05, ** p <.01, *** p <.001.

#### CNNs

In CNNs, depending on the layer and network, only 5-35% of the total units had a slope near 0.5 (Figure 6B-E). This distribution was very different from that of LO where 75% of the voxels had a slope near 0.5. This confirmed the observation made earlier when we inspected the distribution of the unit responses (see Figure 2A-B) and once again showed that extracting the slope across all the units could significantly obscure differences in the overall unit response distributions. For the CNN units that were near 0.5, we did see improvement in pattern averaging, with 10/14 (71.4%) of the layers that did not previously show strong pattern averaging now showing increased averaging regardless of image background (see the asterisks marking the significance levels in Figure 6B & 6D). However, only 3/14 (21.4%) of these layers now showed averaging no different from LO. Thus, better averaging in unit response amplitudes improved but rarely led to full pattern averaging similar to that found in the human LO.

### Evaluating the Effect of Shape Training on Unit and Pattern Normalization

Because CNNs trained on ImageNet images are biased toward texture rather than shape processing (Geirhos et al., 2019), we evaluated whether CNNs biased towards shape processing may exhibit more LO like averaging in both response amplitude and response patterns. We tested 3 versions of Resnet-50 trained with stylized ImageNet images that emphasizes shape processing: Resnet-50 trained on Stylized ImageNet alone (RN50-SIN), Resnet-50 trained on both Original and Stylized ImageNet, and Resnet-50 trained on both Original and Stylized ImageNet (RN50-SININ), with fine-tuning on ImageNet (RN50-SININ-IN) (Geirhos et al., 2019).

At the single unit level, when we compared the slopes of Resnet-50 trained on just the Original ImageNet images with those that were trained on Stylized ImageNet images, we saw the biggest improvement occurred in RN50-SININ-IN (Figure 3C). The original Resnet-50 was only comparable to LO in the final layer, but RN50-SININ-IN was comparable in the last two layers. Meanwhile, at the population level, such shape-training did not improve averaging in response patterns (Figure 4C). Thus, training Resnet-50 to emphasize shape rather than texture processing showed some improvement at the single unit level, but not at the population level.

## Discussion

Previous monkey neurophysiology, human fMRI and CNN studies have reported that neural/CNN responses to a pair of unrelated objects can be well predicted by the average responses to each constituent object shown in isolation, allowing primate vision to recover the identity of an object regardless of whether or not other objects are encoded concurrently. Such an averaging relationship has been found at the single unit level in the slope of response amplitudes of macaque IT neurons and CNN units to paired and single objects (e.g., Reynolds et al., 1999; Zoccolan et al., 2005; Jacob et al., 2021) and at the population level in response patterns of fMRI voxels in the human ventral object processing region (e.g., MacEvoy & Epstein, 2009 & 2011; Reddy, Kanwisher & VanRullen, 2009; Jeong and Xu, 2017). Although it is assumed that averaging at the single unit and population levels must reflect the same underlying process, this assumption, however, has never been directly verified. By evaluating the slope of single unit responses, object-categorization-trained CNN units are also shown to exhibit the same averaging responses as the macaque IT neurons (Jacob et al., 2021). However, averaging the slope across units ignores the overall response distribution, potentially obscuring important differences in how CNNs and the primate brain represent multiple objects. To address these questions, here we first tested whether averaging at the single unit level similar to that found in IT neurons may exist in single fMRI voxels. Conversely, we tested whether averaging in response patterns as observed in human fMRI studies may be present in CNN unit response patterns. We then examined whether voxels/units showing better averaging in response amplitude exhibit better response pattern averaging at the population level. To accomplish these goals, we analyzed fMRI responses from the human LO from an existing data set (Jeong & Xu, 2017). We also examined responses from five CNNs pretrained for object categorization, including Alexnet, Cornet-S, Googlenet, Resnet-50, and VGG-19.

In human LO, replicating Jeong and Xu (2017), we observed averaging in fMRI response pattern between paired and single objects. More strikingly, we also observed averaging in the response amplitude of single fMRI voxels similar to that reported in single neuron recordings in macaque IT studies (Zoccolan et al., 2005). Specifically, across all the included fMRI voxels, the slope of the linear regression between an fMRI voxel’s response to a pair of objects and its summed response to the single objects was indistinguishable from that of the IT neurons. The same results were obtained whether the top 75 most reliable voxels or all the voxels were included, demonstrating the robustness of the effect. To our knowledge, such a result has not been reported before. In a previous study, MacEvoy and Epstein (2009) documented the slope of individual fMRI voxels but did not assess the slope across all the fMRI voxels. Here by directly applying the method used in neurophysiology, we show that the same result reported in neurophysiology studies is present in fMRI voxels even though fMRI is an indirect measure of neural activity with each voxel containing millions of neurons. Our finding further demonstrates the existence of functional smoothness in neuronal representation previously argued to enable the success of fMRI to capture brain functions (Guest & Love, 2017). At the single-neuron level, averaging is argued to be employed to avoid response saturation which would result in the loss of object identity when multiple objects are encoded together (MacEvoy and Epstein, 2009). It is quite remarkable that such a neuronal response property is preserved in fMRI voxel responses. Further analyses revealed that about 75% of LO voxels showed a slope close to 0.5 when the slope was measured within each voxel. More importantly, voxels showing better averaging in response amplitude (i.e., with a slope close to 0.5) also exhibited better averaging in response patterns. We thus verified for the first time a direct connection between averaging at the single unit (i.e., voxel) level and that at the population level.

CNNs have been considered by some as the current best models of the primate ventral visual system (Cichy & Kaiser, 2019; Kubilius et al., 2019). Meanwhile, large discrepancies in object representation also exist between the primate brain and CNNs (e.g., Geirhos et al., 2019; Serre, 2019; Xu & Vaziri-Pashkam, 2021a, 2021b, 2022). In a previous study, Jacob et al. (2021) observed averaging in unit responses in the higher layers of VGG-16 pretrained for object categorization. Here in the higher layers of five CNNs with varying architecture, processing depth, and the presence/absence of recurrent processing, we observed in about half of the sample layers response amplitude averaging in the CNN units comparable to those found in the human LO (i.e., the slope across all the included units did not differ from that of LO). Except for Googlenet, which showed equally good averaging as the human LO in all the sampled layers, for the other four CNNs, there was a general increase in averaging from the lower to the higher sampled layers. Overall, these results replicated and extended the results of Jacob et al. (2021) and showed that averaging, as assessed by the response amplitude slope across all the units, exists in CNN unit responses for representing a pair of objects at the higher levels of CNN visual processing.

Despite the prevalence of averaging in CNN single unit responses, except for the final layers of Resnet-50, no CNN higher layers showed averaging in response patterns comparable to that of the human LO regardless of differences in the exact architecture, depth, and presence/absence of recurrent processing. For example, the recurrent CNN we tested, Cornet-S, which was designed to closely model ventral visual processing, did not outperform the other CNNs. This was also true for layers showing good correspondence with LO in object representational structure as reported in a separate study (Xu & Vaziri-Pashkam, 2021a; note that the final sampled layer of Resnet-50, which showed the best pattern averaging as LO, did not show the best correspondence with LO in representational structure). There is thus a discrepancy between unit and pattern responses in CNNs. Further investigation revealed that, in many of the higher CNN layers sampled, the distribution of unit responses to paired and single objects did not resemble that of the human LO or macaque IT. In fact, when we divided up the units into two groups based on whether the response amplitude of a unit showed good or bad averaging (i.e., whether the slope was near or far from 0.5), we found that only a minority of the CNN units showed good averaging (5%-35% depending on the layer) compared to the majority of the LO voxels that showed good averaging (75%). Thus, extracting the slope across all the units and comparing that with the brain obscured significant distributional differences between the brain and CNNs, leading to the impression that CNNs represent paired objects in a similar way as the primate brain, but in fact the majority do not. Like the LO voxels, CNN units with good averaging also tended to show better averaging in response patterns, but this rarely led to full pattern averaging similar to that found in the human LO.

In the present study, we also considered three factors that may impact the results. In the fMRI experiment, objects were presented within a white rectangle on a gray background. It is possible that this gray background could affect object averaging as CNNs wouldn’t “filter out” the gray background as human participants would. We thus examined CNN responses to objects on both the gray and white backgrounds, but found very little effect of background on CNN unit response amplitude or response pattern. In the main analyses, we only considered a simple average in which the response of each object contributed equally to the prediction of the response of the paired objects. In additional analyses, we also weighed the two objects differently but found that an equal weighting best predicted the response of object pairs from single objects in both the human LO and higher CNN layers. Lastly, to account for the texture bias observed in CNNs, we examined responses from Resnet-50 trained with the stylized versions of ImageNet images that emphasized shape processing (Geirhos et al., 2019). While we observed some improvement in averaging in single unit responses, we saw no improvement in pattern averaging. Such a CNN training regime thus does not appear to make the CNN representation of the paired objects to be more brain-like.

Overall, we found that averaging existed in both single fMRI voxels and voxel population responses in the human LO, with better averaging at the single voxels leading to better averaging in fMRI response patterns. Although CNNs exhibited averaging at the unit level when the slope of response amplitudes for single and pair objects were considered across all the units, detailed investigation revealed that unit response distribution in most cases did not resemble those of the human LO or macaque IT. Consequently, averaging at the population level for the CNN units did not match that of LO. The whole is thus not equal to the average of its parts in CNNs. This indicates the existence of some interactions between the individual objects in a pair that are not present in the human brain, making the identities of the individual objects variable in CNNs at higher levels of visual processing when they are encoded together with other objects.

Such a discrepancy from primate vision is notable, as one of the hallmarks of primate high-level vision is its ability to extract object identity features among changes in non-identity features and form transformation-tolerant object representations (DiCarlo and Cox, 2007; DiCarlo et al., 2012; Tacchetti et al., 2018). This allows us to rapidly recognize an object under different viewing conditions. One aspect of this is clutter tolerance, the ability to recover the identity of an object regardless of the encoding context (Rust & DiCarlo, 2010). The lack of brain-like averaging for representing multiple visual objects in CNNs as shown in the present study suggests that CNNs do not form the same kind of transformation-tolerance visual object representations as the human brain. This echoes the finding from another recent study which shows that, unlike human vision, CNNs do not maintain the object representational structure at higher levels of visual processing when objects undergo transformations involving changes such as position or size (Xu & Vaziri-Pashkam, 2022).

In the primate brain, neural averaging has been linked to a normalization process that involves dividing the response of a neuron by a factor that includes a weighted sum of the activity of a pool of neurons through feedforward, lateral or feedback connections (Carandini & Heeger, 2012; Reynolds & Heeger, 2009; Heeger, 1992). Future CNN architecture development may explicitly impose such a circuit property at higher levels of visual processing. It may also be possible to impose brain-like object averaging in CNN unit responses within the current architecture during training. Such manipulations may not only make CNNs exhibit brain-like averaging in object representations but could also potentially improve other aspects of their performance, making CNNs both better models for the primate brain and better models for object recognition.

## Materials

In this study we reanalyzed data from an existing fMRI data set (Jeong & Xu, 2017) where participants viewed both object pairs and their constituent single objects in a 1-back repetition detection task. We extracted fMRI responses from lateral occipital cortex (LO), a higher ventral visual object processing region homologous to the macaque IT. We also extracted CNN unit responses to the same images from five CNNs pre-trained on object recognition using ImageNet images (Deng et al., 2009). We examined both shallower networks, including Alexnet (Krizhevsky et al., 2012) and VGG-19 (Simonyan & Zisserman, 2014), and deeper networks, including Googlenet (Szegedy et al., 2015) and Resnet-50 (He et al., 2016). We also included a recurrent network, Cornet-S, that has been shown to capture the recurrent processing in macaque IT cortex with a shallower structure, and is argued to be one of the current best models of the primate ventral visual system (Kar et al., 2019; Kubilius et al., 2019). As an additional test, because CNNs trained on ImageNet images are biased toward texture rather than shape processing, we evaluated Resnet-50 trained with stylized ImageNet images that emphasized shape processing (Geirhos et al., 2019). We examined single unit responses in human LO fMRI voxels and in CNN higher layer units. We also examined population responses in human LO fMRI voxel response patterns and in CNN unit response patterns. We then compared single unit and population responses both within human LO and CNNs and between the two systems.

The details of the fMRI experiment has been reported in a previous publication (Jeong & Xu, 2017). They are summarized here for the readers’ convenience.

### Participants

Ten paid participants (eight women) took part in the study with informed consent. They all had normal or corrected-to-normal visual acuity, were right-handed, and between 18 and 35 years old (M = 29.33 years, SD = 3.08 years). One additional participant was tested but excluded from data analysis due to excessive head motion (>3 mm). The study was approved by the institutional review board of Harvard University.

### Experimental Design and Procedures

#### Main Experiment

Participants viewed blocks of images in the main experiment (Figure 1). Each single-object block contained a sequential presentation of 10 images from the same object category either all above or all below the central fixation. Each object pair block contained a sequential presentation of two streams of 10 exemplars from two different categories with one always above and one always below the central fixation. Participants performed a 1-back repetition detection task and pressed a response key whenever they detected an immediate repetition of the exemplar. A repetition occurred twice in each block. In object pair blocks, the repetition occurred randomly in either the upper or lower location. The presentation order of the stimulus blocks and the presentation order of the exemplars within each block were randomly chosen.

Four object categories (shoe, bike, guitar, and couch) with each containing 10 different exemplars were shown. All the exemplars from a given object category were shown in the same view and thus shared a similar outline. This allowed us to increase the difficulty of the 1-back task, thereby increasing participants’ attentional engagement on the task. This also resulted in objects from different categories to be more distinctive from each other. There were eight unique single-object blocks (4 object categories × 2 locations) and 12 unique object pair blocks (with all possible combinations of object categories and locations included).

Each exemplar image subtended approximately 5.5° × 2.8°. Two white square placeholders (7° × 4.7°), marking the two exemplar locations, were shown above and below the central fixation throughout a block of trials. The distance between the central fixation and the center of each placeholder was 3.2°. Each stimulus block lasted 8 sec and contained 10 images, with each appearing 300 msec followed by a 500-msec blank interval. Fixation blocks, which lasted 8 sec, were inserted at the beginning and end of the run and between each stimulus block. Each run contained 20 stimulus blocks, with each unique stimulus block appearing once, and 21 fixation blocks. Each participant was tested with 10 runs, each lasting 5 min 36 sec. Participants’ eye movements during the main experiment were monitored with an EyeLink 1000 eye tracker to ensure proper central fixation.

#### LO Localizer

Following Xu and Jeong (2015), to localize LO, participants viewed blocks of sequentially presented object and noise images (both subtended approximately 12° × 12°). Each object image contained four unique objects shown above, below, and to the left and right of the central fixation (the distance between the fixation and the center of each object was 4°). Gray-scaled photographs of everyday objects were used as the object stimuli. To prevent grouping between the objects, objects appeared on white placeholders (4.5° × 3.6°) that were visible throughout an object image block. Noise images were generated by phase-scrambling the object images used. Each block lasted 16 sec and contained 20 images, with each image appearing for 500 msec followed by a 300-msec blank display. Participants were asked to detect the direction of a slight spatial jitter (either horizontal or vertical), which occurred randomly once in every 10 images. Eight object blocks and eight noise blocks were included in each run. Each participant was tested with two or three runs, each lasting 4 min 40 sec.

### MRI Methods

fMRI data were acquired from a Siemens (Erlangen, Germany) Tim Trio 3-T scanner at the Harvard Center for Brain Science (Cambridge, MA). Participants viewed images back-projected onto a screen at the rear of the scanner bore through an angled mirror mounted on the head coil. All experiments were controlled by an Apple MacBook Pro laptop running MATLAB (The MathWorks Natick, MA) with Psychtoolbox extensions (Brainard, 1997). For the anatomical images, high-resolution T1-weighted images were acquired (repetition time = 2200 msec, echo time = 1.54 msec, flip angle = 7°, 144 slices, matrix size = 256 × 256, and voxel size = 1 × 1 × 1 mm). Functional data in the main experiment and in the LO localizer were acquired using the same gradient-echo echo-planar T2*-weighted sequence (repetition time = 2000 msec, echo time = 30 msec, flip angle = 90°, 31 slices, matrix size = 72 × 72, voxel size = 3 × 3 × 3 mm, 168 volumes for the main experiment and 140 volumes for the LO localizer).

### Data Analysis

#### fMRI data processing

fMRI data were analyzed using FreeSurfer (surfer.nmr.mgh.harvard.edu), FsFast (Dale et al. 1999), and in-house MATLAB and Python codes. FMRI data preprocessing included 3D motion correction, slice timing correction and linear and quadratic trend removal. No smoothing was applied to the data. The first two volumes of all functional runs were also discarded. All the analysis for the main experiment was performed in the volume. The ROIs were selected on the surface and then projected back to the volume for further analysis.

LO was defined separately in each participant as a cluster of continuous voxels in the lateral occipital cortex showing higher activations to the object images than to the noise images (*p* < .001 uncorrected; Figure 1B). For two participants, the threshold of *p* < .001 resulted in too few voxels so the threshold was relaxed to *p* < .01.

For each participant, a general linear model with 20 factors (8 single-object conditions and 12 object-pair conditions) was applied to the fMRI data from the main experiment, and beta values were extracted from each stimulus block in each run and in each voxel of LO.

To equate the number of voxels across participants and to increase power, we selected 75 most reliable voxels in each ROI using reliability-based voxel selection (Tarhan & Konkle, 2020). This is based on the fact that, across the participants, the LO ROI ranged from 75 to 240 voxels before voxel selection. This method selects voxels whose response profiles are consistent across odd and even halves of the runs and works well when there are around 15 conditions. To implement this method, for each voxel, we calculated the split-half reliability by first averaging the runs within the odd and even halves and then correlating the resulting averaged responses for all conditions (20 in total) across the even and odd halves. We then selected the top 75 voxels with the highest correlations. The 75 voxels chosen had a moderate split-half reliability with an average of around *r = 0.20* across the voxels and across the participants. We also analyze the data including all the voxels in LO and found virtually identical results.

#### CNN details

We tested 5 CNNs in our analyses (See Table 1). We examined both shallower networks, including Alexnet (Krizhevsky et al., 2012) and VGG-19 (Simonyan & Zisserman, 2014), and deeper networks, including Googlenet (Szegedy et al., 2015) and Resnet-50 (He et al., 2016). We also included a recurrent network, Cornet-S, that has been shown to capture the recurrent processing in macaque IT cortex with a shallower structure and argued to be one of the current best models of the primate ventral visual system (Kar et al., 2019; Kubilius et al., 2019). All the CNNs used were pretrained with ImageNet images. As an additional test, because CNNs trained on ImageNet images are biased toward texture rather than shape processing, we evaluated 3 versions of Resnet-50 trained with stylized ImageNet images that emphasizes shape processing: Resnet-50 trained on Stylized ImageNet alone (RN50-SIN), Resnet-50 trained on both Original and Stylized ImageNet (RN50-SININ), and Resnet-50 trained on both Original and Stylized ImageNet, with fine-tuning on ImageNet (RN50-SININ-IN)(see Geirhos et al., 2019).

**Table 1.** CNNs and layers examined. All of the layers shown were examined in previous studies and are representative of the higher levels visual processing hierarchy (Xu & Vaziri-Pashkam, 2021a, 2021b & 2022; Mocz et al., 2022). Here, for each CNN, following Jacob et al., (2021), we only examined normalization in the higher-level layers, as only these layers show a large portion of units that respond to objects in both locations. Additionally, bold layers were those that showed best correspondence with LO as shown by Xu & Vaziri-Pashkam (2021a).

| CNN name | Depth/Blocks | Layers | N of Total Layers Sampled | Sampled Layer Names and Locations (indicated in the parenthesis) |
| --- | --- | --- | --- | --- |
| Alexnet | 8 | 25 | 3 | **'fc6'** (17), **'fc7'** (20), 'fc8' (23) |
| Cornet-S | 4 | 42 | 3 | 'IT_output' - IT (38), 'decoder_avgpool' - avgpool (39), 'decoder_output' - output (42) |
| Googlenet | 22 | 144 | 3 | **'pool4-3x3_s2'** - **pool4**(111), **'pool5-7x7_s1'** - **pool5**(140), 'loss3-classifier' - classifier (142) |
| Resnet-50 | 50 | 177 | 3 | **'activation_40_relu'** - **act40** (141), **'avg_pool'** - **avgpool** (174), 'fc1000' (175) |
| VGG-19 | 19 | 47 | 3 | **'fc6'** (39), **'fc7'** (42), 'fc8' (45) |

In previous studies (Mocz et al., 2022; Tang et al., 2022; Xu & Vaziri-Pashkam, 2021a, & 2021b & 2022), we sampled 6 to 8 pooling and fully connected (FC) layers representative of all stages of visual processing in these same CNNs and compared them with the human ventral visual processing areas. Following Jacob et al., (2021) who have previously examined averaging in CNN single units, here we included only the 3 highest layers we sampled previously, as only these layers contained a large portion of units that respond to objects at both locations (see Table 1 for the specific CNN layers sampled). We included pooling layers here because they typically mark the end of processing for a block of layers when information is pooled to be passed on to the next block of layers. When there were no obvious pooling layers present, the last layer of a block was chosen. It has been shown previously that such a sampling procedure captures the evolution of the representation trajectory fairly well, if not fully, as adjacent layers exhibit identical or very similar representations (Taylor & Xu, 2021). In each CNN, the layer(s) showing stronger correlation with LO in their representational structure for real-world objects than other layers tested in the same CNN (see Vaziri-Pashkam & Xu, 2021a) were marked with bold font in all the results figures. For a given CNN layer, we extracted CNN unit responses to the 20 possible image conditions that the human participants saw in the scanner. These images all showed objects within a white rectangle on a gray background. As human participants could ignore the gray background and focus on the objects only but CNNs could not do so, we also extracted CNN unit responses to the same images on white backgrounds to assess the effect of background color on response averaging. Cornet-S and the three Stylized versions of Resnet-50 were implemented in Python. All other CNNs were implemented in Matlab. The output from all CNNs was analyzed and compared with brain responses using Python and R.

### Evaluating the Unit Response to Single and Paired Objects

We followed the method of Zoccolan et al. (2005) and Jacob et al. (2021) to assess unit response to single and paired objects in single fMRI voxels in human LO and in CNN units (Figure 1C). In LO, for each participant, we first averaged the beta weights across all the runs for each condition in each voxel. We then extracted paired and single object responses from each of the 12 object pairs and averaged across all the pairs to generate an averaged paired and single object responses. We included all the LO voxels for a given participant and extracted the slope of the linear regression between a voxel’s response to a pair of objects and its summed response to the corresponding single objects shown in isolation. The resulting slope was then averaged across the participants. The average slope should be 0.5 if the single voxel response to an object pair can be perfectly predicted by the average response of the corresponding objects shown alone. In a previous single-cell analysis of monkey IT, a slope of 0.55 was reported (Zoccolan et al., 2005). We thus compared our slope results to both 0.5 and 0.55 as baselines. For each CNN layer examined, we only included in the regression analysis units that were responsive to objects at both locations; in other words, units that showed a non-zero variance across the four single object conditions at each of the two locations. We then compared the slopes of the linear regressions of the CNN layers exacted using the same procedure as outlined above with that of human LO.

### Evaluating the Pattern Response to Single and Paired Objects

We followed the method of Jeong & Xu (2017) to assess pattern response to single and paired objects in fMRI response patterns in human LO and in CNN unit response patterns (Figure 1d). In LO, to extract voxel response patterns, following established practice (e.g., Kamitani & Tong, 2006), for each participant, we first z-normalized the beta values across all voxels for each condition in each run to remove amplitude differences between conditions and runs. We then divided the data into odd and even runs and averaged the runs within each half. We performed two correlations: correlating the voxel response pattern of the same object pair between odd and even runs (actual pair) and correlating the voxel response of an object pair and the average of its constituent objects shown alone between odd and even runs (averaged pair). To account for the fMRI measurement noise, we derived a noise-corrected correlation by dividing average pair correlation with actual pair correlation. If the noise-corrected correlation is no different from 1, it would indicate that the representation of an object pair can be perfectly predicted by the average representation of its constituent objects shown in isolation. This was done for each object pair and the results were averaged across all the object pairs for each participant. For CNNs, because there was no noise in the data, we simply calculated the response pattern correlation of an object pair and the average of its constituent objects. Similar to the CNN unit response analysis, we only included units that were responsive to objects at both locations. We then directly compared whether the correlations from the CNN layers were significantly different from that of the human LO.

To examine whether mechanisms other than a simple average or sum can predict the representation of an object pair, we additionally tested in both human LO and CNNs whether or not patterns generated by a weighted average model would better predict that of the actual object pair. To do so, we systematically varied the contribution of the two isolated objects’ patterns from 0% to 100%, with 10% increments in between. The resulting weighted average was then evaluated in the same way as before.

### Evaluating the Relationship between Unit Response and Pattern Response

Here we examined whether fMRI voxels/CNN units with better averaging in response amplitude would exhibit better averaging in response pattern. To do so, we extracted the slope of the linear regression between an fMRI voxel/unit’s response to a pair of objects and its summed response to the corresponding single objects shown in isolation as before. We then sorted the voxels/CNN units into two groups based on whether their slopes were close to 0.5, defined as having a slope between 0.45 and 0.55 (referred to as the *near* condition), or whether they were far from 0.5, defined as having a slope less than 0.45 or greater than 0.55 (referred to as the *far* condition). Finally, we evaluated averaging in response patterns for the near and far voxels/units separately, and tested whether the near ones showed better pattern averaging than the far ones. We also noted the percentage of voxels/units that were either near or far.

#### Experimental Design and Statistical Analyses

Ten human participants took part in the experiment. The factors described in the previous methods sections were evaluated at the group level using t-tests. One-tailed t tests were performed when the comparison in one direction was meaningful. We corrected for multiple comparisons in all post-hoc analyses using the Benjamini-Hochberg method (Benjamini & Hochberg, 1995). For the fMRI analyses of LO, we corrected for two comparisons with a baseline level of normalization (0.5 or 0.55 for the unit response analysis and 1 for the population response analysis), accounting for including either all of the voxels or the top 75 most reliable voxels. For both the CNN unit and population analyses, for each CNN, we corrected for six comparisons with LO (three layers each and two background colors). When examining the relationship between CNN unit and population analyses, for each CNN, and for each background color, we corrected for nine comparisons (three layers and three groups of units included, i.e., near, far, and all units). We also calculated effect size using Cohen’s D (Cohen, 1969; Cohen, 1988). All of the above statistical tests were conducted using R (R Core Team, 2018).

## Author contributions

VM, MC and YX conceived the study; SKJ and YX designed the fMRI experiments; SKJ implemented and conducted the experiments, and collected fMRI data; VM performed fMRI data preprocessing and beta weights extraction, and performed the normalization analysis; VM and YX wrote the manuscript with comments from SKJ and MC.

## Acknowledgements

We thank members of Visual Cognitive Neuroscience Lab, Turk-Browne Lab, Holmes Lab, Yale Cognitive and Neural Computation Lab, and Ilker Yildirim for their helpful feedback on this project. This research was supported by the National Institute of Health Grants 1R01EY030854 and 1R01EY022355 to Y.X..

## Data availability statement

Data for the present study will be posted shortly on the osf website.

## References

Bao P., & Tsao, D. Y. (2018) Representation of multiple objects in macaque category-selective areas. Nature Communications, 9, 1774.

Benjamini, Y., & Hochberg, Y. (1995). Controlling the False Discovery Rate: A Practical and Powerful Approach to Multiple Testing. Journal of the Royal Statistical Society. Series B (Methodological), 57(1), 289–300. JSTOR.

Brainard, D. H. (1997). The Psychophysics Toolbox. Spatial Vision, 10(4), 433–436. https://doi.org/10.1163/156856897X00357

Carandini, M., & Heeger, D. J. (2012). Normalization as a canonical neural computation. Nature Reviews Neuroscience, 13(1), 51–62. https://doi.org/10.1038/nrn3136

Cichy, R. M., & Kaiser, D. (2019). Deep Neural Networks as Scientific Models. Trends in Cognitive Sciences, 23(4), 305–317. https://doi.org/10.1016/j.tics.2019.01.009

Cichy, R. M., Khosla, A., Pantazis, D., Torralba, A., & Oliva, A. (2016). Comparison of deep neural networks to spatio-temporal cortical dynamics of human visual object recognition reveals hierarchical correspondence. Scientific Reports, 6(1), 27755. https://doi.org/10.1038/srep27755

Cohen, J. (1969). Statistical power analysis for the behavioral sciences. New York: Academic Press.

Cohen, J. (1988). Statistical power analysis for the behavioral sciences (2nd ed.). Hillsdale, NJ: Erlbaum.

Dale, A. M., Fischl, B., & Sereno, M. I. (1999). Cortical Surface-Based Analysis: I. Segmentation and Surface Reconstruction. NeuroImage, 9(2), 179–194. https://doi.org/10.1006/nimg.1998.0395

Deng, J., Dong, W., Socher, R., Li, L.-J., Li, K., & Fei-Fei, L. (2009). ImageNet: A large-scale hierarchical image database. 2009 IEEE Conference on Computer Vision and Pattern Recognition, 248–255. https://doi.org/10.1109/CVPR.2009.5206848

DiCarlo, J. J., & Cox, D. D. (2007). Untangling invariant object recognition. Trends in Cognitive Science, 11, 333–341.

DiCarlo, J. J., Zoccolan, D., & Rust, R. C. (2012). How does the brain solve visual object recognition? Neuron 73, 415–434.

Eickenberg, M., Gramfort, A., Varoquaux, G., & Thirion, B. (2017). Seeing it all: Convolutional network layers map the function of the human visual system. NeuroImage, 152, 184– 194. https://doi.org/10.1016/j.neuroimage.2016.10.001

Geirhos, R., Rubisch, P., Michaelis, C., Bethge, M., Wichmann, F. A., & Brendel, W. (2019). ImageNet-trained CNNs are biased towards texture; increasing shape bias improves accuracy and robustness. ArXiv:1811.12231 [Cs, q-Bio, Stat]. http://arxiv.org/abs/1811.12231

Güçlü, U., & van Gerven, M. A. J. (2015). Deep Neural Networks Reveal a Gradient in the Complexity of Neural Representations across the Ventral Stream. Journal of Neuroscience, 35(27), 10005–10014. https://doi.org/10.1523/JNEUROSCI.5023-14.2015

Guest, O., & Love, B. C. (2017). What the success of brain imaging implies about the neural code. ELife, 6, e21397. https://doi.org/10.7554/eLife.21397

He, K., Zhang, X., Ren, S., & Sun, J. (2016). Deep Residual Learning for Image Recognition. 2016 IEEE Conference on Computer Vision and Pattern Recognition (CVPR), 770–778. https://doi.org/10.1109/CVPR.2016.90

Heeger, D. J. (1992). Normalization of cell responses in cat striate cortex. Visual Neuroscience, 9(2), 181–197. https://doi.org/10.1017/S0952523800009640

Jacob, G., Pramod, R. T., Katti, H., & Arun, S. P. (2021). Qualitative similarities and differences in visual object representations between brains and deep networks. Nature Communications, 12(1), 1872. https://doi.org/10.1038/s41467-021-22078-3

Jeong, S. K., & Xu, Y. (2017). Task-context-dependent Linear Representation of Multiple Visual Objects in Human Parietal Cortex. Journal of Cognitive Neuroscience, 29(10), 1778–1789. https://doi.org/10.1162/jocn_a_01156

Kamitani, Y. & Tong, F. (2005). Decoding the visual and subjective contents of the human brain. Nature Neuroscience 8, 679–685.

Kar, K., Kubilius, J., Schmidt, K., Issa, E. B., & DiCarlo, J. J. (2019). Evidence that recurrent circuits are critical to the ventral stream’s execution of core object recognition behavior. Nature Neuroscience, 22(6), 974–983. https://doi.org/10.1038/s41593-019-0392-5

Kay, K. N. (2018). Principles for models of neural information processing. NeuroImage 180, 101– 109.

Khaligh-Razavi, S.-M., & Kriegeskorte, N. (2014). Deep Supervised, but Not Unsupervised, Models May Explain IT Cortical Representation. PLOS Computational Biology, 10(11), e1003915. https://doi.org/10.1371/journal.pcbi.1003915

Kietzmann, T. C., Spoerer, C. J., Sörensen, L. K. A., Cichy, R. M., Hauk, O., & Kriegeskorte, N. (2019). Recurrence is required to capture the representational dynamics of the human visual system. Proceedings of the National Academy of Sciences, 116(43), 21854–21863. https://doi.org/10.1073/pnas.1905544116

Kliger, L., & Yovel, G. (2020). The functional organization of high-level visual cortex determines the representation of complex visual stimuli. Journal of Neuroscience, 40, 7545–7558.

Kriegeskorte, N. (2015). Deep neural networks: a new framework for modeling biological vision and brain information processing. Annual review of vision science, 1, 417–446.

Krizhevsky, A., Sutskever, I., & Hinton, G. E. (2012). ImageNet classification with deep convolutional neural networks. Advances in neural information processing systems, 25, 1097–1105.

Kubilius J, Schrimpf M, Hong H (2019) Brain-like object recognition with high-performing shallow recurrent ANNs. In: NeurIPS | 2019, Thirty-Third Conference on Neural Information Processing Systems. San Diego: Neural Information Processing Systems.

MacEvoy, S. P., & Epstein, R. A. (2009). Decoding the Representation of Multiple Simultaneous Objects in Human Occipitotemporal Cortex. Current Biology, 19(11), 943–947. https://doi.org/10.1016/j.cub.2009.04.020

MacEvoy, S. P., & Epstein, R. A. (2011). Constructing scenes from objects in human occipitotemporal cortex. Nature Neuroscience, 14(10), 1323–1329. https://doi.org/10.1038/nn.2903

Marr, D. (1982). Vision: A Computational Investigation into the Human Representation and Processing of Visual Information. W.H. Freeman, San Francisco, CA.

Mocz, V., Vaziri-Pashkam, M., Chun, M., & Xu, Y. (2022). Predicting identity-preserving object transformations in human posterior parietal cortex and convolutional neural networks. Journal of Cognitive Neuroscience, 34, 2406–2435.

R Core Team (2018) R: a language and environment for statistical computing. Vienna: R Foundation for Statistical Computing. Available at http://www.R-project.org/.

Rajalingham, R., Issa, E. B., Bashivan, P., Kar, K., Schmidt, K., & DiCarlo, J. J. (2018). Large-Scale, High-Resolution Comparison of the Core Visual Object Recognition Behavior of Humans, Monkeys, and State-of-the-Art Deep Artificial Neural Networks. The Journal of Neuroscience, 38(33), 7255–7269. https://doi.org/10.1523/JNEUROSCI.0388-18.2018

Reddy, L. & Kanwisher, N. (2007). Category selectivity in the ventral visual pathway confers robustness to clutter and diverted attention. Current Biology, 17, 2067–2072.

Reddy, L., Kanwisher, N. G., & VanRullen, R. (2009). Attention and biased competition in multi-voxel object representations. Proceedings of the National Academy of Sciences, 106(50), 21447–21452. https://doi.org/10.1073/pnas.0907330106

Rust, N. C., & DiCarlo, J. J. (2010). Selectivity and tolerance (“invariance”) both increase as visual information propagates from cortical area V4 to IT. Journal of Neuroscience, 30, 12978– 12995.

Reynolds, J. H., Chelazzi, L., & Desimone, R. (1999). Competitive Mechanisms Subserve Attention in Macaque Areas V2 and V4. Journal of Neuroscience, 19(5), 1736–1753. https://doi.org/10.1523/JNEUROSCI.19-05-01736.1999

Reynolds, J. H., & Heeger, D. J. (2009). The Normalization Model of Attention. Neuron, 61(2), 168–185. https://doi.org/10.1016/j.neuron.2009.01.002

Serre, T. (2019). Deep Learning: The Good, the Bad, and the Ugly. Annual Review of Vision Science, 5(1), 399–426. https://doi.org/10.1146/annurev-vision-091718-014951

Simonyan, K., & Zisserman, A. (2014). Very Deep Convolutional Networks for Large-Scale Image Recognition. ArXiv:1409.1556 [Cs]. http://arxiv.org/abs/1409.1556

Szegedy, C., Wei Liu, Yangqing Jia, Sermanet, P., Reed, S., Anguelov, D., Erhan, D., Vanhoucke, V., & Rabinovich, A. (2015). Going deeper with convolutions. 2015 IEEE Conference on Computer Vision and Pattern Recognition (CVPR), 1–9. https://doi.org/10.1109/CVPR.2015.7298594

Tacchetti, A., Isik, L., & Poggio, T. A. (2018). Invariant recognition shapes neural representations of visual input. Annual Review of Vision Science, 4, 403–422.

Tang, K., Chin, M., Chun, M., & Xu, Y. (2022). The contribution of object identity and configuration to scene representation in convolutional neural networks. PLoS ONE, 17, e0270667.

Tarhan, L., & Konkle, T. (2019). Reliability-based voxel selection. NeuroImage, 116350. https://doi.org/10.1016/j.neuroimage.2019.116350

Taylor, J., & Xu, Y. (2021). Conjunctive Coding of Color and Shape in Convolutional Neural Networks. Journal of Vision, 20(11), 400–400.

Xu, Y., & Vaziri-Pashkam, M. (2021a). Limits to visual representational correspondence between convolutional neural networks and the human brain. Nature Communications, 12(1), 2065. https://doi.org/10.1038/s41467-021-22244-7

Xu, Y., & Vaziri-Pashkam, M. (2021b). Examining the Coding Strength of Object Identity and Nonidentity Features in Human Occipito-Temporal Cortex and Convolutional Neural Networks. The Journal of Neuroscience, 41(19), 4234–4252. https://doi.org/10.1523/JNEUROSCI.1993-20.2021

Xu, Y. & Vaziri-Pashkam, M. (2022). Understanding transformation tolerant visual object representations in the human brain and convolutional neural networks. Neuroimage, 263, 119635.

Yamins, D. L. K., Hong, H., Cadieu, C. F., Solomon, E. A., Seibert, D., & DiCarlo, J. J. (2014). Performance-optimized hierarchical models predict neural responses in higher visual cortex. Proceedings of the National Academy of Sciences, 111(23), 8619–8624. https://doi.org/10.1073/pnas.1403112111

Yamins, Daniel L. K., & DiCarlo, J. J. (2016). Using goal-driven deep learning models to understand sensory cortex. Nature Neuroscience, 19(3), 356–365. https://doi.org/10.1038/nn.4244

Zoccolan, D. (2005). Multiple Object Response Normalization in Monkey Inferotemporal Cortex. Journal of Neuroscience, 25(36), 8150–8164. https://doi.org/10.1523/JNEUROSCI.2058-05.2005

